# Persistent Mullerian duct syndrome in dogs – a new insight into organization of *AMH* and *AMHR2* genes

**DOI:** 10.1101/2024.11.28.625841

**Authors:** Paulina Krzeminska

## Abstract

Persistent Müllerian Duct Syndrome (PMDS) is a rare congenital disorder in males, characterized by the presence of Müllerian duct derivatives despite normal testes and external genitalia. This condition is typically linked to a dysfunction in the anti-Müllerian hormone (AMH) or its receptor (AMHR2), both of which are critical for the regression of the Müllerian ducts. In dogs, PMDS is particularly frequent in the Miniature Schnauzer breed, although cases have also been reported in other breeds, such as the Yorkshire Terrier. To date, a single causative variant has been identified in the *AMHR2* gene, but only in Miniature Schnauzers. No deleterious variants have been found in the *AMH* gene; however, with the exception of one report, most studies have not sequenced the entire exon 5.

This study provides novel insights into the genomic organization of canine *AMH* and *AMHR2* genes through bioinformatics and in silico analyses of previously reported whole-genome sequencing (WGS) data from a Yorkshire Terrier affected by PMDS. The results indicate that current canine genome assemblies (ROS_Cfam_1.0; CanFam4, and CanFam6) contain a complete reference sequence for the *AMH* gene, unlike the earlier CanFam3.1 genome. However, next-generation sequencing technologies (WGS and RNA-seq) face challenges due to technical limitations in analyzing GC-rich repetitive elements present in exon 5 of canine *AMH* gene. In contrast, the genomic structure of the *AMHR2* gene remains inaccurately represented in the current ROS_Cfam_1.0 genome (eight instead of eleven exons), while both CanFam4 and CanFam6 contain additional and unknown nucleotide/amino acid sequences. The CanFam3.1 genome assembly still provides the most accurate annotation for canine *AMHR2* gene.

Based on these findings, re-sequencing of the *AMH* gene in previously reported dogs affected by PMDS using the methodology proposed in recent literature is recommended. Further attention should be given to comparative analyses to assess whether the dog’s genome contains accurate information about genes or proteins that correspond to human orthologs.

## INTRODUCTION

Persistent Müllerian Duct Syndrome (PMDS) is a rare congenital disorder of sexual development in individuals with a male karyotype (XY) (Picard and Josso, 2019). PMDS is characterized by the presence of the Müllerian duct derivatives, such as the uterus and fallopian tubes, which typically develop in females but should regress in males during fetal development due to the action of anti-Mullerian hormone (AMH), secreted by Sertoli cells (Brunello and Rey, 2022).

AMH, a member of the Transforming Growth Factor-beta family, binds to its primary receptor (AMHR2), which then recruits a type I receptor (Howard et al., 2022). AMHR2 is expressed in the Mullerian ducts, and the AMH-AMHR2 signaling pathway is crucial for the regression their regression (Cate, 2022). It is important to note that AMH is initially produced as a pre-pro-peptide, with its mature chain located at the C-terminal end of the protein. The dimerization of two AMH molecules, followed by the cleavage between the propeptide and the mature chain, is essential for activation of the AMH-AMHR2 signaling pathway (Cate, 2022). However, the mature chains, actinh as homodimers, remain associated with the prodomains via non-covalent bonds (Josso and Picard, 2022). Multiple isoforms of the AMH protein have been identified; however bioative isoforms must contain the C-terminal mature chain (McLennan and Pankhurst, 2015).

Cases of PMDS typically present as phenotypic males with developed testes. However, the majority of them exhibit cryptorchidism, and when both gonads are undescended, patients are stertile (Picard et al., 2017). Cryptorchidism is also observed in dogs with PMDS (Meyers-Wallen, 2012). It is important to note that cases with descended testes or unilateral cryptorchidism can still pass a causative variant to their offspring due to active spermatogenesis. In humans, a comprehensive review of reported PMDS cases revealed that around 19% of affected males were fertile and able to father at least one child (Mullen et al., 2019; Picard et al., 2017). In dogs, approximately 50% of PMDS cases had descended testes, or at least one testis, with active spermatogenesis (Meyers-Wallen, 2012; Wu et al., 2009). For this reason, understanding the genetic etiology of PMDS is particularly crucial for limiting the population spread of the pathogenic variants.

PMDS is a well-documented condition in humans, with approximately 200 cases reported, typically caused by mutations in the *AMHR2* or *AMH* genes (Brunello and Rey, 2022). Mutations in the *AMH* and *AMHR2* genes have been identified in the majority of coding exons, including numerous mutations in exon 5 of the *AMH*, which encodes the bioactive chain (Josso et al., 2013). However, around 10% of male cases are classified as idiopathic, with an unknown genetic cause (Picard and Josso, 2019). PMDS has also been observed in domestic animals, particularly in dogs (**Supplementary Table 1**), in cats (Meyers-Wallen, 2012; Rozynek et al., 2024), and in a single goat (Haibel and Rojko, 1990). In contrast, no PMDS cases have been reported in horses, cattle, pigs, or sheep.

Molecular analyses of two candidate genes (*AMH* and *AMHR2*) have been carried out in only a few dogs and a single cat. In dogs, a causative mutation in the *AMHR2* gene has been identified exclusively in the Miniature Schnauzer breed, confirming a sex-limited autosomal recessive inheritance model (Wu et al., 2009). On the other hand, no mutations in the *AMHR2* or *AMH* genes were found in Basset Hounds [Pop et al., 2017], a Belgian Malinois (Smit et al., 2018), a German Shepherd (De Lorenzi et al., 2018), or a Yorkshire Terrier (Nowacka- Woszuk et al., 2022). PMDS cases have also been reported in Cocker Spaniel and Pomeranian dogs; however, molecular analyses of both genes were not performed (Cinti et al., 2021; Vignoli et al., 2020). In a single European shorthair cat with PMDS, both genes were re-sequenced, but no deleterious DNA variants were found, even though the genes exhibited polymorphic variations (Rozynek et al., 2024).

The Miniature Schnauzer breed has been recognized as one of the dog breeds most affected by PMDS, with numerous reports of affected cases published (Breshears and Peters, 2011; Brown et al., 1976; Marshall et al., 1982; Vegter et al., 2010). Since the testes of these dogs produced active AMH protein, researchers focused on analyzing the *AMHR2* gene, where a causative mutation was identified in the North American population of this breed (Wu et al., 2009). A rapid molecular test was subsequently proposed to identify this single substitution in exon 3 of the *AMHR2* gene (Pujar and Meyers-Wallen, 2009).

In recent years, addidional cases of PMDS have been reported in Miniature Schnauzer dogs from Europe and South America, as well as a new case from the USA, in which the mutation in the *AMHR2* gene was also detected (Dzimira et al., 2018; Nogueira et al., 2019; Welsh et al., 2023).

The high frequency of PMDS in Miniature Schnauzers, particularly in North America, indicates that selective breeding practices may have contributed to the spread of a harmful DNA variant in exon 3 of the *AMHR2* gene within this population. Research showed that in a group of 216 Miniature Schnautzers, the frequency of a deleterious allele was estimated at 16%, with heterozygous carriers accounting for 27% of the population (Smit et al., 2018).

Given this data, it is essential for breeders to genotype all Miniature Schnautzers for this DNA variant. This proactive approach will help identify and eliminate dogs affected by PMDS, as well as healthy carriers of the deleterious variant, from breeding programs. This strategy could contribute to improving the health of future generations of the Miniature Schnauzer population. However, several previously reported PMDS cases suggest that other DNA variants may be responsible for abnormal sexual development in other breeds.

It was assumed that the unsuccessful search for causative variants in both candidate genes (*AMH* and *AMHR2*) may have been caused by limited knowledge of their organization in the dog genome. Given that the previous canine genome assembly (CanFam3.1) was incomplete, the aim of this study was to perform new bioinformatic and *in silico* analyses of both genes, which were previously studied in a PMDS dog by Nowacka-Woszuk et al. (2022).

## MATERIAL

The raw whole-genome sequencing (WGS) data for five male Yorkshire Terrier dogs, studied by Nowacka-Woszuk et al. (2022) and deposited in the the NCBI Sequence Read Archive (SRA) database (accession number: PRJNA779963), were downloaded.

In addition, raw RNA sequencing (RNA-seq) data from four control dogs studied by Stachowiak et al. (2024) and deposited in the NCBI Sequence Read Archive (accession number: PRJNA901164) were also used.

## METHODS

The downloaded WGS data were mapped using the Burrows-Wheeler Aligner (BWA) (version 0.7.17-r1188) (Li and Durbin, 2009) to four canine genome assemblies:

- the previous **CanFam3.1** (accession: GCF_000002285.3),
- the **ROS_Cfam_1.0** (accession: GCF_014441545.1),
- the UU_Cfam_GSD_1.0 (accession: GCF_011100685.1, **CanFam4**),
- the Dog10K_Boxer_Tasha (accession: GCF_000002285.5, **CanFam6**).

The quality of the mapped BAM files was assessed using the Qualimap tool (version v2.2.1) (Okonechnikov et al., 2016). Samtools (version 1.13; htslib 1.13+ds) (Li et al., 2009) was used to sort (samtools sort) and index (samtools index) of BAM files. The obtained mapping results were visualized using the Integrative Genomics Viewer (IGV) (Robinson et al., 2011).

The results from the most recent RNA-seq analysis were aligned to the Cfam_ROS_1.0 genome assembly using the Rsubread package in RStudio (v4.4.1) (Liao et al., 2019; R Core Team, 2023). Sorting and indexing were performed with Samtools, as described above.

## IN SILICO ANALYSES

DNA sequences of the *AMH* gene were downloaded from the NBCI database for six species: **mouse** (NM_007445.3), **bovine** (NM_173890.1), **pig** (XM_021081532.1), **human** (NM_000479.5), **dog** (NM_001314127.1), and **cat** (XM_011288073.4).

Protein sequences for the AMH and AMHR2 were obtained from the Uniprot database: **mouse** (P27106 and Q8K592), **bovine** (P03972 and E1BHR7), **pig** (P79295 and A0A4X1WCQ3), **human** (P03971 and Q16671), **dog** (A0A8C0N2S3 and A0A8C0TD47), and **cat** (A0A2I2V0G5 and M3W4X5).

In addition, protein sequence for canine AMHR2 was obtained from the Ensembl archive (ENSCAFP00000052775.1; CanFam3.1) and the NCBI database (XP_038293864.1 for ROS_Cfam_1.0; XP_038433716.1 for CanFam4; XP_543632.4 for CanFam6).

A GC content calculator (https://en.vectorbuilder.com/tool/gc-content-calculator.html) was used to compare the sequences of exon 5 of the *AMH* gene between selected species.

DNA sequences of exon 5 of the *AMH* gene for the selected species were examined for the presence of tandem repeats using the Tandem Repeats Finder tool (https://tandem.bu.edu/trf/basic_submit) (Benson, 1999).

Multiple alignments of DNA sequences for exon 5 of the *AMH* gene were performed using the Clustal Omega tool (https://www.ebi.ac.uk/jdispatcher/msa/clustalo?stype=dna). The same tool was used for aligning the protein sequences.

Protein structure modeling was performed using the Swiss-Model tool (https://swissmodel.expasy.org/) (Waterhouse et al., 2018). In addition, PyMOL software (DeLano, 2002) was applied to mark domains in the predicted protein models with selected colors.

Figures dispaying sequencing data were generated using screenshots from the Integrative Genomics Viewer software. These images were subsequently modified to condense and highlight relevant information.

## RESULTS

The raw sequencing data from five male Yorkshire Terrier dogs were successfully aligned to the *Canis lupus familiaris* genome assemblies: ROS_Cfam_1.0, CanFam3.1, CanFam4, and CanFam6. A summary of the quality statistics for case 6942 is presented in **Supplementary Table 2**. Among these genome assemblies, **ROS_Cfam_1.0** and **CanFam4** show the highest mapping rates (99.43% and 99.44%, respectively). The **ROS_Cfam_1.0** exhibits high mapping quality (42.68) and good mean coverage with low standard deviation. **CanFam4** has a similar mapping rate but lower mapping quality (16.15) and moderate coverage. **CanFam6** achieves the highest mean mapping quality (49.53) and mean coverage (44.96), but also has a higher duplication rate (20.85%). In contrast, the older **CanFam3.1** assembly has lower mapping quality and more variable coverage.

The PMDS dog referred to as case 6942 by Nowacka-Woszuk et al. (2022) was the only male dog with a homozygous CC genotype in the 3’UTR of the *HSD3B2* gene (rs851059986). Based on this genotype, the raw sequencing data from WGS for case 6942 were identified (**Supplementary Figure 1**) and further analyzed in detail.

### AMH analysis

Based on the the current genome assemblies (ROS_Cfam_1.0, CanFam4, and CanFam6), visualization of the BAM files revealed deep coverage for five exons of the *AMH* gene, except for a large fragment of exon 5 (Figure 1A **and Supplementary Figure 2**). Due to the absence of reads within exon 5 of the *AMH* gene, a multiple nucleotide sequence alignment was performed across selected species (Figure 2A). This analysis identified additional nucleotide sequences in dogs and cats, predominantly composed of G and C nucleotides (highlighted in green and yellow). A critical nucleotide sequence for AMH activity, marked in purple, showed low conservation, primarily due to differences in the mouse sequence.

**Figure 1.**
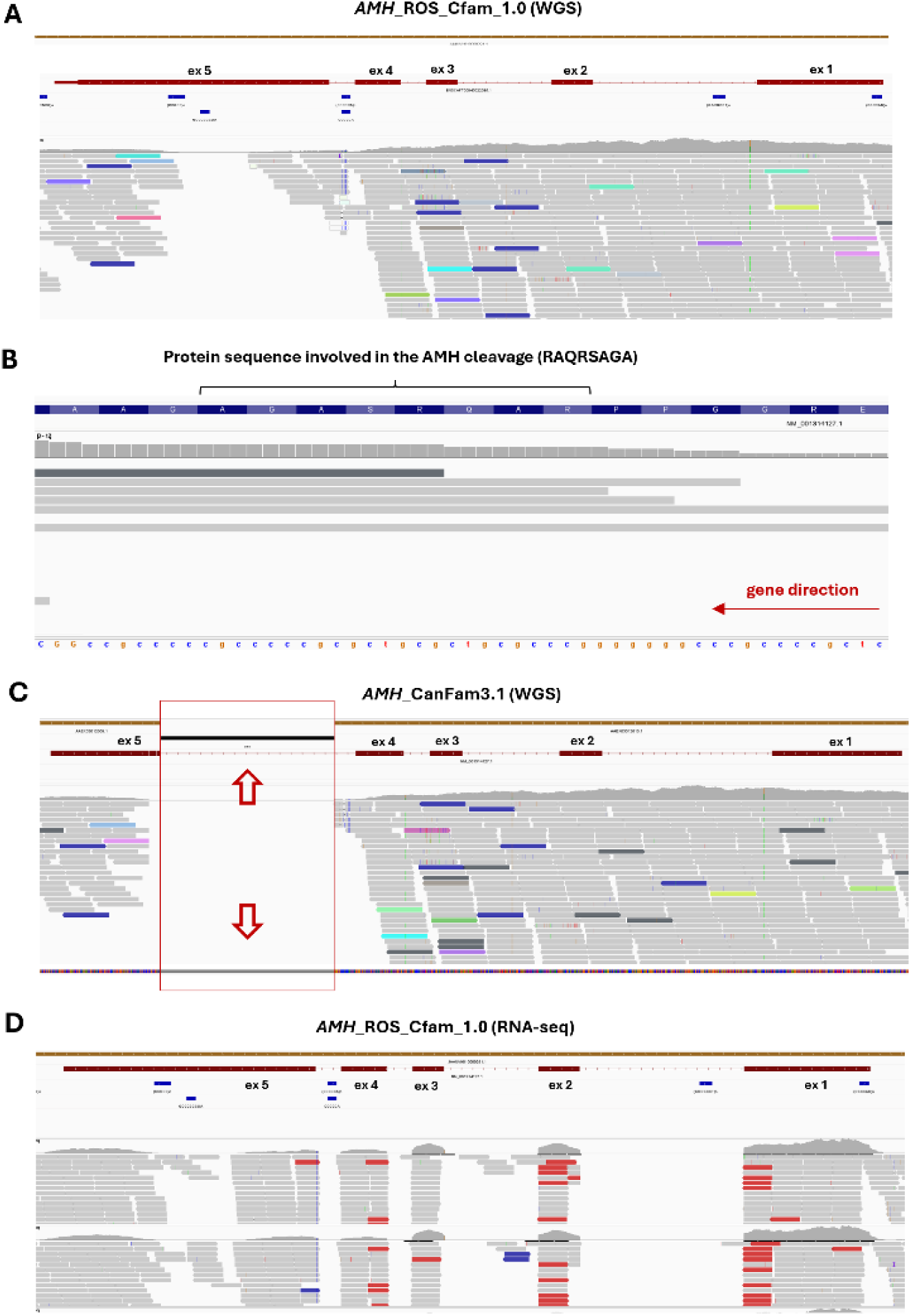
Results of sequencing read mapping for the *AMH* gene. **A)** WGS data mapped to the ROS_Cfam_1.0 assembly, showing a large gap of coverage within exon 5. **B)** Indication of the fragment encoding an important RAQRSAGA peptide, with very low coverage. **C)** WGS data mapped to the CanFam3.1 assembly, indicating a gap in the reference sequence (two red arrows). **D)** RNA-seq data mapped to the ROS_Cfam_1.0 assembly, showing poor coverage wihin exon 5.

**Figure 2.**
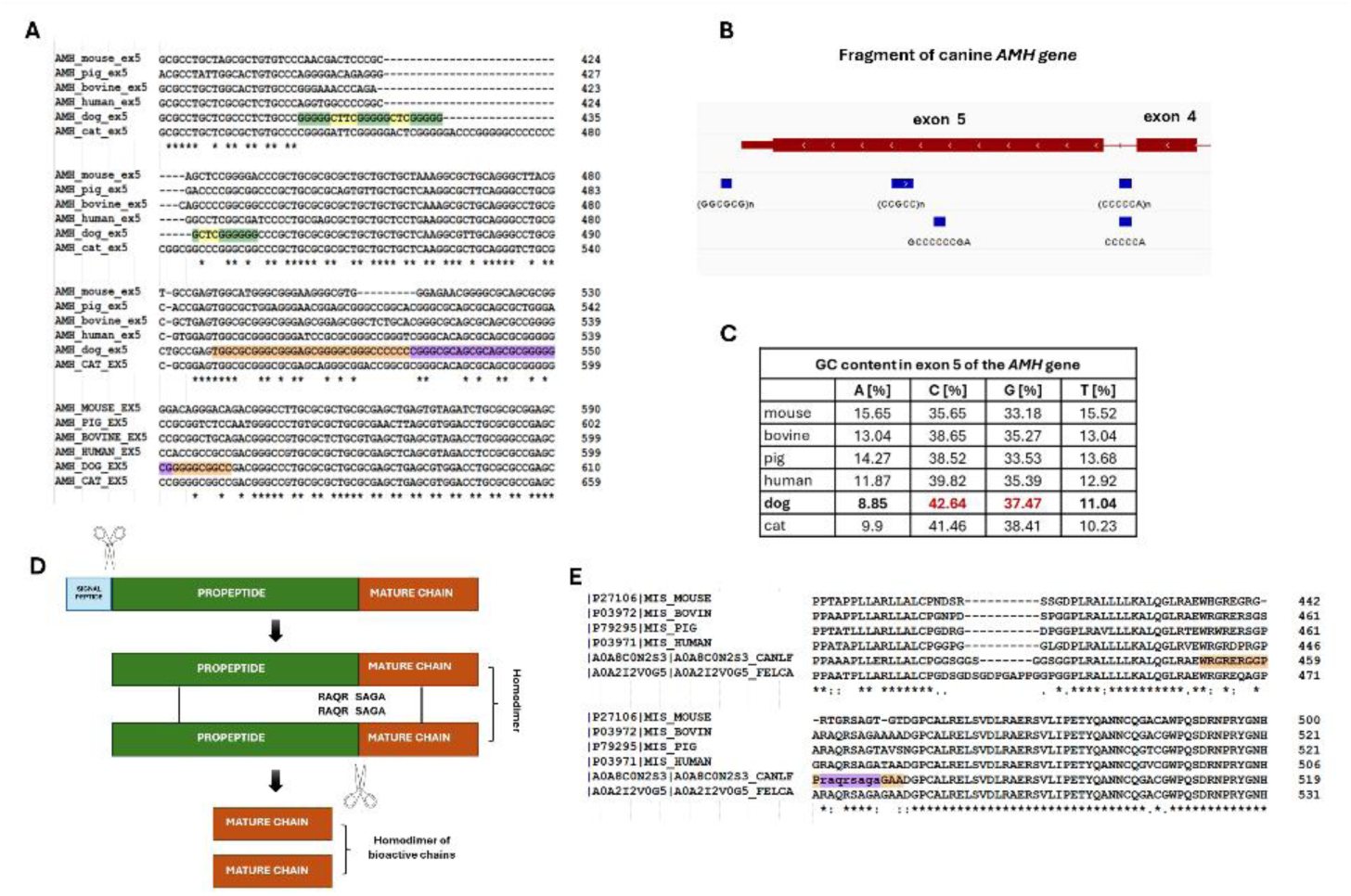
In silico analyses of the *AMH* gene. A) Nucleotide sequence comparison for exon 5 of the *AMH* gene across selected species. B) Presence of repetitive elements within exon 5, analyzed using the IGV tool. C) Comparison of nucleotide content within exon 5 for selected species. D) AMH protein domains and processing from inactive to active mature chains. E) Comparison of the C-terminal AMH protein sequence for selected species, highlighing essential sequence for AMH processing (marked in purple).

Furthermore, analysis using the Tandem Repeats Finder tool showed that dog has GC-rich tandem repeats in this region (Figure 2A), a finding also confirmed using the IGV tool (Figure 2B). A GC content analysis across species demonstrated that all species have a GC-rich exon 5 (Figure 2C). Therefore, the previously reported challenges in analyzing canine exon 5 are likely due not only to the GC-rich sequence but also the presence of GC-rich tandem repeats.

Exon 5 of the *AMH* gene encodes the C-terminal region of the AMH protein, which is crucial for the dimerization of its mature chains (Figure 2D). This region contains the RAQRSAGA polypeptide (Figure 2D **and 2E**), a sequence essential for the proper processing of the AMH protein. In all examined Yorkshire Terriers, the nucleotide sequence encoding the RAQRSAGA amino acids was not fully covered by sequencing reads (Figure 1B).

In contrast, when mapping to the CanFam3.1 genome assembly, it was observed that a large part of exon 5 contained a gap in the assembly, with the sequence in this region represented as “NNNN” (Figure 1C). The Ensembl database versions from 2015 and 2019 did not include the entire canine *AMH* gene, further complicating the analysis of this region (https://www.ensembl.org/info/website/archives/index.html?redirect=no).

RNA-seq data from additional control dogs, mapped to the ROS_Cfam_1.0 genome assembly, indicated that the fragment of exon 5 of the *AMH* gene was also poorly covered by sequencing reads, although some reads were observed mapping to this region (Figure 1D).

### AMHR2 analysis

The *AMHR2* gene in the ROS_Cfam_1.0 genome consists of eight exons, all of which exhibited deep coverage in whole-genome sequencing (WGS) data from five Yorkshire Terrier dogs (Figure 3A) and RNA-seq data from additional control dogs (Figure 3C). However, a small gap in coverage was observed within exon 2 when using both sequencing technologies (WGS and RNA-seq) (Figure 3B).

**Figure 3.**
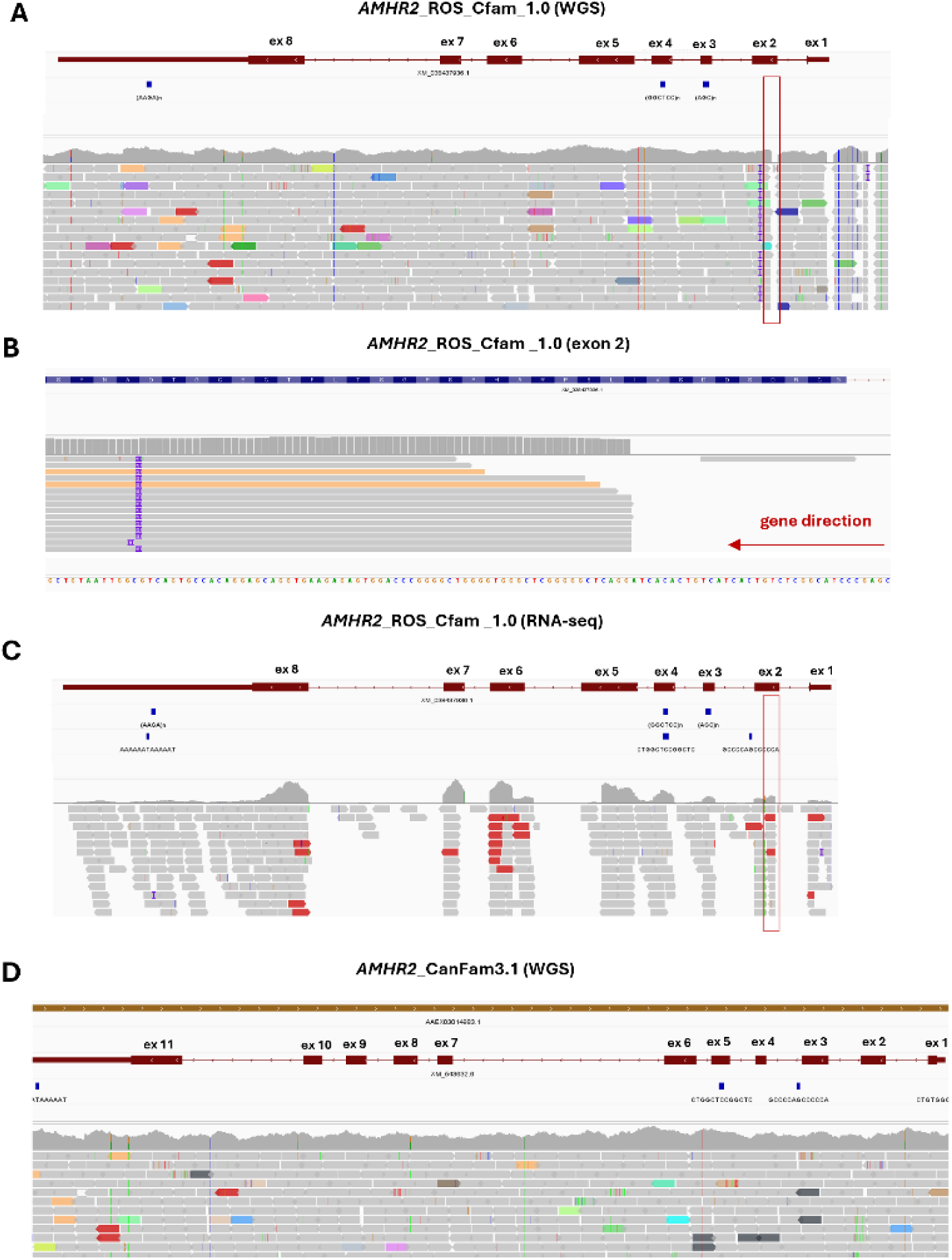
Results of sequencing read mapping for the *AMHR2* gene. **A)** WGS data mapped to the ROS_Cfam_1.0 assembly, showing eight coding exons and a small coverage gap within exon 2. **B)** Indication of the fragment encoded by the sequence without read coverage. **C)** RNA-seq data mapped to the ROS_Cfam_1.0 assembly, indicating a small coverage gap within exon 2. **D)** WGS data mapped to the CanFam3.1 assembly, showing 11 coding exons with complete and deep read coverage.

In contrast, mapping to the CanFam3.1 (Figure 3D), CanFam4, and CanFam6 genome assemblies (**Supplementary Figure 3**) revealed that the canine *AMHR2* gene contains 11 exons, all of which were fully covered by sequencing reads. This exon count is consistent with that found in other species included in the comparative analysis (data not shown).

To further investigate, amino acids sequences for selected species were obtained from the Uniprot database and aligned, revealing shorter sequences in dog and cat compared to other species. Specifically, the protein sequence consists of 573 amino acids in human, 477 amino acids in dog, and 483 amino acids in cat (**Supplementary Figure 4**). To evaluate how different versions of the canine genome predict the AMHR2 protein sequence, comparison and structural prediction analyses were performed for the following:

- AMHR2_cf_ROS_Cfam_1.0 (XP_038293864.1),
- AMHR2_cf_CanFam3.1 ( ENSCAFP00000052775.1; the Ensembl database from 2019),
- AMHR2_cf_CanFam4 (XP_038433716.1),
- AMHR2_cf_CanFam6 (XP_543632.4),
- AMHR2_human (Q16671).

It was noted that the AMHR2_ROS_Cfam_1.0, AMHR2_CanFam4, and AMHR2_CanFam6 amino acid sequences contain an „X”, at the C-terminal end instead of a proper amino acid abbreviation. To predict the protein structure, this „X” was replaced with glycine (G), a small and neutral amino acid. The alignment comparison is presented in **Supplementary Figure 5**, while the predicted structures for all sequences are shown in **Suplementary Figure 6.**

The AMHR2_ROS_Cfam_1.0 variant (497 amino acids) was shorter than the other proteins (CanFam3.1: 565 aa; both CanFam4 and CanFam6: 612 aa; human: 573 aa). The N-terminal region, corresponding to the extracellular domain (marked in yellow), was absent in AMHR2_ROS_Cfam_1.0 compared to CanFam3.1, the CanFam4/CanFam6 (which are identical), and human AMHR2. In addition, several gaps were observed within the AMHR2_ROS_Cfam_1.0 protein. The transmembrane domain (marked in green) was conserved across all proteins. However, the C-terminal intracellular domain of AMHR2_ROS_Cfam_1.0, AMHR2_CanFam4, and the AMHR2_CanFam6 was longer and did not correspond to human AMHR2 or AMHR2_CanFam3.1 sequences. Moreover, these additional polipeptides were not found in any of the templates used by the Swiss Model tool during structural prediction.

## DISCUSSION

A case of a single **Yorkshire Terrier** dog with normal external genitalia, including a fully developed penis and testes, but with a uterus, was reported by Nowacka-Woszuk et al. (2022). This phenotype suggested that the dog was affected by Persistent Müllerian Duct Syndrome (PMDS). The authors conducted whole-genome sequencing (WGS) analysis, but no deleterious variants were identified in either the *AMH* or *AMHR2* genes. It should be noted that the raw sequencing data were mapped to the canine genome assembly CanFam3.1, which had an incomplete representation of the *AMH* gene, specifically excluding exon 5.

Given that previous canine genome assembly (CanFam3.1) was incomplete for the *AMH* gene, this study aimed to perform new bioinformatics and in silico analyses on the previously described case, focusing on both the *AMH* and *AMHR2* genes. The findings provide significant insights into the structure of the *AMH* and *AMHR2* genes and their protein sequences in dogs, particularly in relations to PMDS.

Visualization of BAM files from both WGS and RNA-seq revealed deep coverage of the *AMH* gene. However, a notable gap was identified in exon 5, which encodes a critical C-terminal region of the AMH protein, including the essential RAQRSAGA polypeptide sequence. This suggests that case #6942 may harbor a deleterious variant in this exon. Thus, further sequence analysis of this case is strongly recommended.

The lack of full coverage in this region of the analyzed Yorkshire Terriers raises concerns about the adequacy and effectiveness of next- generation sequencing techniques for GC-rich regions. These findings highlight the limitations of such technologies in analyzing challenging genomic regions. The presence of GC-rich tandem repeats, as identified by both the Tandem Repeats Finder and IGV tools, further complicates the sequencing of the canine AMH gene.

It is well known that GC-rich sequences are difficult to amplify using polymerase chain reaction (PCR), as both the template DNA and the primers often form secondary structures, which inhibit amplification. The most common recommendation is to add DMSO and betaine to the PCR mixture to prevent the formation of these secondary structures (Green and Sambrook, 2019). Another key approach is „slowdown” PCR, where the annnealing temperature is gradually decreased to improve amplification (Frey et al., 2008). Recently, a new class of oligonucleotide reagents, known as disruptors, has been introduced. These reagents are specifically designed to counteract problematic secondary structures, providing an effective solution for amplifying particularly difficult sequence that are resistant to DMSO and betaine additives (Ma and Zheng, 2023).

A Belgian Malinois dog with PMDS was the first case in which both the *AMH* and *AMHR2* genes were fully analyzed (Smit et al., 2018). The authors of that study were the first to compare the available data for the canine *AMH* gene with that of other species and noted that that coding sequence of the canine *AMH* gene, according to the CanFam3.1 genome assembly from NCBI database, was incomplete. As a result, a set of multiple primers, along with modified PCR conditions (including DMSO and betaine additives, as well as a decrease in annealing temperature by one-third of a degree Celsius per cycle ), were used to amplify and sequence exon 5 of the canine *AMH* gene. However, no causative DNA variant in the *AMH* gene was identified in the PMDS Belgian Malinois. Nevertheless, Smit et al. (2018) proposed a methodology that enables the amplification and Sanger sequencing of exon 5 of the canine *AMH* gene. This approach should be considered for all dogs suspected of PMDS with an unknown genetic background, including previously reported cases (**Supplementary Table 1**). Specifically, a Basset Hound dog should be of interest, as it demonstrated low AMH activity [(Pop et al., 2017; Pop et al., 2015). In contrast, both the Yorkshire Terrier (Szabo et al., 2023) and German Shepherd (De Lorenzi et al., 2018) had an ovotestis and ovarian tissue, respectively. The presence of ovarian tissue suggests that these dogs exhibit a more complex disorder of sexual development, including abnormal gonadal development. Since these dogs had an 78,XY karyotype, a potential deleterious variant may be localized in genes involved in testicular differentiation, such as the *SOX9*, *FGF9*, or *NR5A1* genes.

It is also worth noting that several genes involved in mammalian sexual development exhibit high GC content (results observed in this analysis, although not shown). One example is the *INSL3* gene, which shows low and inadequate coverage in the promoter region. This gene plays a key role in testicular descent and has been previously analyzed in cryptorchid dogs (Krzeminska et al., 2022). Sanger sequencing of exon 1 of this gene was challenging due to the presence of a CpG island that spans the entire exon. Another example is the *NR5A1* gene, for which no reads were found due to GC-rich tandem repeats within the 5’ region of the gene. The human *NR5A1* gene consists of seven coding exons and encodes the SF-1 factor, which is involved in both gonadal development and steroidogenesis (Elzaiat et al., 2022; Köhler and Achermann, 2010). Numerous mutations have been identified in human *NR5A1* gene in patients with various DSD phenotype (Domenice et al., 2016). In a single dog, a heterozygous large deletion spanning exons 1-4 was reported (Nowacka-Woszuk et al., 2020). That study included four coding exons of the canine *NR5A1*. One of the available versions of this gene annotation at that time, ENSCFA00000032206.3, described five exons. The current ROS_Cfam_1.0 assembly identifies seven coding exons for canine *NR5A1* gene. An intriguing example is the *FOXL2* gene, which demonstrated very low coverage in its two exons, likely due to numerous GC-rich repeats. The *FOXL2* gene is crucial for ovarian development and was previously analyzed in 78,XX DSD dogs with testes or ovotestes (Salamon et al., 2015). However, the authors discontinued sequencing exon 1 of the *FOXL2* gene due to the technical difficulties. In addition, many 78,XX DSD cases were included in a whole-genome sequencing (WGS) analysis (Nowacka-Woszuk et al., 2022). As mentioned, this gene showed very low coverage in the WGS data. Therefore, it seems worthwhile to re- analyze these dogs using Sanger sequencing and the PCR conditions proposed by Smit et al. (2018). The final gene, *SOX9*, which is crucial for testicular development, also exhibited several GC-rich repeats, including one in exon 1, where very low coverage was observed.

Moreover, this analysis also revealed the presence of additional nucleotide sequences, predominantly composed of G and C nucleotides, in the coding sequences of the *AMH* gene in both dog and cats. This raises questions about the mechanism behind their presence in the canine and feline genomes, as well as the function of the addidional PTAAA and DSGDPGAPPG amino acid sequences found in cats.

In contrast, the analysis of the *AMHR2* gene demonstrated robust coverage across all eight exons, based on the current genome assembly for dogs (ROS_Cfam_1.0). However, a gap in mapping within exon 2 was observed in both whole-genome sequencing (WGS) and RNA-seq data, with only eight exons identified in this assembly. Notably, other species included in the *in silico* analysis contain 11 exons.

Mapping reads to other genome assemblies (CanFam3.1, CanFam4, and CanFam6) revealed the expected number of exon (11), and demonstrated deep and complete coverage by sequencing reads. This analysis showed that the canine genome still contains incorrect gene annotations, despite extensive efforts to sequence and describe it in details.

In addition, amino acid sequence alignment revealed differences in protein length between dog and human, with the canine AMHR2 sequence being shorter based on the ROS_Cfam_1.0 genome. This difference is due to the absence of the N-terminal region of the protein, as well as the presence of several gaps within the sequence. In contrast, the length of canine AMHR2 based on the CanFam4 and CanFam6 genomes is similar to that of human protein. However, all three of these current canine genome versions contain an error („X”) in the amino acid sequence, as well as a longer C-terminal end of the protein that does not correspond to human sequence. The most similar sequence and structure were found for AMHR2 in the CanFam3.1 genome assembly. It appears that the most accurate gene annotation for canine AMHR2 protein remains the one in the CanFam3.1 genome.

One example of a canine gene with noted differences in exon number is the *ESR1* gene. Pathirana et al. (2010) analyzed this gene in relation to canine cryptorchidism, including 17 exons of the canine *ESR1*. However, both the NCBI database and the Ensembl database currently report only eight coding exons, along with several non-coding. Notably, no isoform of this gene contains 17 exons. The second example is the aforementioned *NR5A1* gene. Given these discrepancies, the search for causative DNA variants in crucial genes for sexual development should be preceded by a comprehensive and comparative analysis of gene organization in other species, primarily humans.

This study demonstrated that the CanFam6 assembly offers the highest mean mapping quality (49.53) and coverage (44.96) with a relatively lower standard deviation among the four assemblies. However, it also showed that gene annotation for the *AMHR2* gene, is not accurate. A recent study presented a French Bulldog dog with abnormal genitalia and performed comprehensive analyses, including whole-genome sequencing (de Gennaro et al., 2024). It is worth noting that the authors used both the CanFam4 and CanFam6 genome assemblies and highlighted that the dog genome still requires a more accurate and detailed representation.

## CONCLUSIONS

Persistent Mullerian Duct Syndrome (PMDS) has been documented across various dog breeds; however, the genetic basis of this condition remains largely unexplored. To date, a causative DNA variant for canine PMDS has only been identified in the *AMHR2* gene, and only in the Miniature Schnauzer breed. Despite the challenges associated with amplifying exon 5 of the *AMH* gene, this region warrants further investiagation, particularly given that causative DNA variants in human PMDS cases are frequently located in exon 5. Therefore, this study strongly recommends re-sequencing the *AMH* in dogs affected by PMDS, following the methodology outlined by Smit et al. (2018). Enhanced investigation of the canine *AMH* gene, particularly through targeted re-sequencing, could provide crucial genetic insights into PMDS across different breeds, advancing both diagnosis and efforts to limit the spread of this syndrome in dogs.

In addition, this study underscores the importance of comparative genomic analyses in uncovering the genetic basis of canine disorders of sexual development (DSD), such as PMDS. The observed variability in *AMHR2* gene organization across different canine genome assemblies highlights the need for in-depth sequence analysis, especially when whole-genome sequencing (WGS) is applied. Furthermore, the coverage gap in the *AMH* gene demonstrates a significant limitation of current sequencing technologies in GC-rich regions with repetitive elements. Since GC-rich repetitive elements are present in several canine genes involved in sexual development, targeted re-sequencing and optimized methodologies could improve our ability to identify causative variants of canine DSDs.

## Author Contributions

PK – designed the study, analyzed the data, and wrote the manuscript.

## References

Benson, G., 1999. Tandem repeats finder: a program to analyze DNA sequences. Nucleic Acids Res 27, 573–580.

Breshears, M.A., Peters, J.L., 2011. Ambiguous genitalia in a fertile, unilaterally cryptorchid male miniature schnauzer dog. Vet Pathol 48, 1038–1040.

Brown, T.T., Burek, J.D., McEntee, K., 1976. Male pseudohermaphroditism, cryptorchism, and Sertoli cell neoplasia in three miniature Schnauzers. J Am Vet Med Assoc 169, 821–825.

Brunello, F.G., Rey, R.A., 2022. AMH and AMHR2 Involvement in Congenital Disorders of Sex Development. Sex Dev 16, 138–146.

Cate, R.L., 2022. Anti-Müllerian Hormone Signal Transduction involved in Müllerian Duct Regression. Front Endocrinol (Lausanne) 13, 905324.

Cinti, F., Sainato, D., Charlesworth, T., 2021. A case of persistent Mullerian duct syndrome in a dog. J Small Anim Pract 62, 311.

de Gennaro, L., Burgio, M., Lacalandra, G.M., Petronella, F., L’Abbate, A., Ravasini, F., Trombetta, B., Rizzo, A., Ventura, M., Cicirelli, V., 2024. Genomic Sequencing to Detect Cross-Breeding Quality in Dogs: An Example Studying Disorders in Sexual Development. International journal of molecular sciences 25.

De Lorenzi, L., Arrighi, S., Groppetti, D., Bonacina, S., Parma, P., 2018. Persistent Müllerian Duct Syndrome in a German Shepherd Dog. Sex Dev 12, 288–294.

DeLano, W., 2002. The PyMOL Molecular Graphics System. DeLano Scientific.

Domenice, S., Machado, A.Z., Ferreira, F.M., Ferraz-de-Souza, B., Lerario, A.M., Lin, L., Nishi, M.Y., Gomes, N.L., da Silva, T.E., Silva, R.B., Correa, R.V., Montenegro, L.R., Narciso, A., Costa, E.M., Achermann, J.C., Mendonca, B.B., 2016. Wide spectrum of NR5A1-related phenotypes in 46,XY and 46,XX individuals. Birth Defects Res C Embryo Today 108, 309–320.

Dzimira, S., Wydooghe, E., Van Soom, A., Van Brantegem, L., Nowacka-Woszuk, J., Szczerbal, I., Switonski, M., 2018. Sertoli Cell Tumour and Uterine Leiomyoma in Miniature Schnauzer Dogs with Persistent Müllerian Duct Syndrome Caused by Mutation in the AMHR2 Gene. J Comp Pathol 161, 20–24.

Elzaiat, M., McElreavey, K., Bashamboo, A., 2022. Genetics of 46,XY gonadal dysgenesis. Best Pract Res Clin Endocrinol Metab 36, 101633.

Frey, U.H., Bachmann, H.S., Peters, J., Siffert, W., 2008. PCR-amplification of GC-rich regions: ’slowdown PCR’. Nat Protoc 3, 1312–1317.

Green, M.R., Sambrook, J., 2019. Polymerase Chain Reaction (PCR) Amplification of GC-Rich Templates. Cold Spring Harb Protoc 2019.

Haibel, G.K., Rojko, J.L., 1990. Persistent müllerian duct syndrome in a goat. Vet Pathol 27, 135–137.

Howard, J.A., Hart, K.N., Thompson, T.B., 2022. Molecular Mechanisms of AMH Signaling. Front Endocrinol (Lausanne) 13, 927824.

Josso, N., Picard, J.Y., 2022. Genetics of anti-Müllerian hormone and its signaling pathway. Best Pract Res Clin Endocrinol Metab 36, 101634.

Josso, N., Rey, R.A., Picard, J.Y., 2013. Anti-müllerian hormone: a valuable addition to the toolbox of the pediatric endocrinologist. International journal of endocrinology 2013, 674105.

Köhler, B., Achermann, J.C., 2010. Update--steroidogenic factor 1 (SF-1, NR5A1). Minerva Endocrinol 35, 73–86.

Krzeminska, P., Nowak, T., Switonski, M., 2022. Isolated cryptorchidism in dogs is not associated with polymorphisms of the INSL3 and AR candidate genes. Anim Genet 53, 233–235.

Li, H., Durbin, R., 2009. Fast and accurate short read alignment with Burrows-Wheeler transform. Bioinformatics 25, 1754–1760.

Li, H., Handsaker, B., Wysoker, A., Fennell, T., Ruan, J., Homer, N., Marth, G., Abecasis, G., Durbin, R., 2009. The Sequence Alignment/Map format and SAMtools. Bioinformatics 25, 2078–2079.

Liao, Y., Smyth, G.K., Shi, W., 2019. The R package Rsubread is easier, faster, cheaper and better for alignment and quantification of RNA sequencing reads. Nucleic Acids Res 47, e47.

Ma, Y., Zheng, M., 2023. Improved PCR by the Use of Disruptors, a New Class of Oligonucleotide Reagents. Methods Mol Biol 2967, 159–171.

Marshall, L.S., Oehlert, M.L., Haskins, M.E., Selden, J.R., Patterson, D.F., 1982. Persistent Müllerian duct syndrome in miniature schnauzers. J Am Vet Med Assoc 181, 798–801.

McLennan, I.S., Pankhurst, M.W., 2015. Anti-Müllerian hormone is a gonadal cytokine with two circulating forms and cryptic actions. J Endocrinol 226, R45–57.

Meyers-Wallen, V.N., 2012. Gonadal and sex differentiation abnormalities of dogs and cats. Sex Dev 6, 46–60.

Mullen, R.D., Ontiveros, A.E., Moses, M.M., Behringer, R.R., 2019. AMH and AMHR2 mutations: A spectrum of reproductive phenotypes across vertebrate species. Dev Biol 455, 1–9.

Nogueira, D.M., Armada, J.L.A., Penedo, D.M., Tannouz, V.G.S., Meyers-Wallen, V.N., 2019. Persistent Mullerian duct Syndrome in a Brazilian miniature schnauzer dog. An Acad Bras Cienc 91, e20180752.

Nowacka-Woszuk, J., Stachowiak, M., Szczerbal, I., Szydlowski, M., Szabelska- Beresewicz, A., Zyprych-Walczak, J., Krzeminska, P., Nowak, T., Lukomska, A., Ligocka, Z., Biezynski, J., Dzimira, S., Nizanski, W., Switonski, M., 2022. Whole genome sequencing identifies a missense polymorphism in PADI6 associated with testicular/ovotesticular XX disorder of sex development in dogs. Genomics 114, 110389.

Nowacka-Woszuk, J., Szczerbal, I., Stachowiak, M., Dzimira, S., Nizanski, W., Biezynski, J., Nowak, T., Gogulski, M., Switonski, M., 2020. Screening for structural variants of four candidate genes in dogs with disorders of sex development revealed the first case of a large deletion in NR5A1. Anim Reprod Sci 223, 106632.

Okonechnikov, K., Conesa, A., García-Alcalde, F., 2016. Qualimap 2: advanced multi-sample quality control for high-throughput sequencing data. Bioinformatics 32, 292–294.

Pathirana, I.N., Tanaka, K., Kawate, N., Tsuji, M., Kida, K., Hatoya, S., Inaba, T., Tamada, H., 2010. Analysis of single nucleotide polymorphisms in the 3’ region of the estrogen receptor 1 gene in normal and cryptorchid Miniature Dachshunds and Chihuahuas. J Reprod Dev 56, 405–410.

Picard, J.Y., Cate, R.L., Racine, C., Josso, N., 2017. The Persistent Müllerian Duct Syndrome: An Update Based Upon a Personal Experience of 157 Cases. Sex Dev 11, 109–125.

Picard, J.Y., Josso, N., 2019. Persistent Müllerian duct syndrome: an update. Reprod Fertil Dev 31, 1240–1245.

Pop, A., Henegariu, O., Micu, R., Sonea, A., Irimie, A., Henegariu, A., Groza, I., 2017. Hormone receptor type 2 antimüllerian gene role in dogs with persistent Müllerian ducts syndrome. Romanian Biotechnological Letters 22, 13029–13034.

Pop, A., Irimie, A., Riu, O.H., Damian, A., Groza, I., 2015. A study regarding antimullerian factor’s activity in basset hound puppy males younger than 120 days. Scientific Papers: Veterinary Medicine, Timișoara XLVIII (48), 156–160.

Pujar, S., Meyers-Wallen, V.N., 2009. A molecular diagnostic test for persistent Müllerian duct syndrome in miniature schnauzer dogs. Sex Dev 3, 326–328.

R Core Team, 2023. A language and environment for statistical computing. Version 4.4.1. . R Foundation for Statistical Computing, Vienna, Austria.

Robinson, J.T., Thorvaldsdóttir, H., Winckler, W., Guttman, M., Lander, E.S., Getz, G., Mesirov, J.P., 2011. Integrative genomics viewer. Nat Biotechnol 29, 24–26.

Rozynek, J., Nowacka-Woszuk, J., Stachowiak, M., Sowinska, N., Lukomska, A., Gruss, M., Switonski, M., Szczerbal, I., 2024. Lack of causative mutation in the AMH and AMHR2 genes in a cat (38,XY) with persistent Mullerian duct syndrome (PMDS). Reprod Domest Anim 59, e14635.

Salamon, S., Nowacka-Woszuk, J., Switonski, M., 2015. Polymorphism of the CTNNB1 and FOXL2 Genes is not Associated with Canine XX Testicular/Ovotesticular Disorder of Sex Development. Folia Biol (Krakow) 63, 57–62.

Smit, M.M., Ekenstedt, K.J., Minor, K.M., Lim, C.K., Leegwater, P., Furrow, E., 2018. Prevalence of the AMHR2 mutation in Miniature Schnauzers and genetic investigation of a Belgian Malinois with persistent Müllerian duct syndrome. Reprod Domest Anim 53, 371–376.

Stachowiak, M., Nowacka-Woszuk, J., Szabelska-Beresewicz, A., Zyprych-Walczak, J., Krzeminska, P., Sosinski, O., Nowak, T., Switonski, M., 2024. A massive alteration of gene expression in undescended testicles of dogs and the association of KAT6A variants with cryptorchidism. Proc Natl Acad Sci U S A 121, e2312724121.

Szabo, Z., Moser, J., Vincenti, S., 2023. Persistent mullerian duct syndrome in a dog. Schweiz Arch Tierheilkd 165, 189–180.

Vegter, A.R., Kooistra, H.S., van Sluijs, F.J., van Bruggen, L.W., Ijzer, J., Zijlstra, C., Okkens, A.C., 2010. Persistent Mullerian duct syndrome in a Miniature Schnauzer dog with signs of feminization and a Sertoli cell tumour. Reprod Domest Anim 45, 447–452.

Vignoli, M., De Amicis, I., Tamburro, R., Quaglione, G., Salviato, N., Collivignarelli, F., Terragni, R., Pastrolin, S., Marruchella, G., 2020. A Case of Adenocarcinoma of Uterus Masculinus in a Pomeranian Dog. Front Vet Sci 7, 337.

Waterhouse, A., Bertoni, M., Bienert, S., Studer, G., Tauriello, G., Gumienny, R., Heer, F.T., de Beer, T.A.P., Rempfer, C., Bordoli, L., Lepore, R., Schwede, T., 2018. SWISS-MODEL: homology modelling of protein structures and complexes. Nucleic Acids Res 46, W296–w303.

Welsh, P.J., McDaniel, K., Goldsmith, E.W., Ramsay, J.D., Conley, A., Owen, T.J., Ambrosini, Y.M., Ciccarelli, M., 2023. Case report: Persistent Müllerian duct syndrome and enlarged prostatic utricle in a male dog. Front Vet Sci 10, 1185621.

Wu, X., Wan, S., Pujar, S., Haskins, M.E., Schlafer, D.H., Lee, M.M., Meyers-Wallen, V.N., 2009. A single base pair mutation encoding a premature stop codon in the MIS type II receptor is responsible for canine persistent Müllerian duct syndrome. J Androl 30, 46–56.

