## Supplementary Materials for "Persistent Mullerian duct syndrome in dogs – a new insight into organization of *AMH* and *AMHR2* genes"

**Krzeminska Paulina**

**Supplementary Table 1.** Summary of canine PMDS cases, including phenotypic features and genetic investigations.

| **Breed** | **No** | **Uterus** | **Other female organs** | **Gonads** | **Cryptorchidism** | **Other male organs** | ***AMH*** | ***AMHR2*** | **Genetic cause** | **Reference** |
| --- | --- | --- | --- | --- | --- | --- | --- | --- | --- | --- |
| **Miniature Schnautzer** | 3 | **yes** | na | testes | bilateral abdominal | na | no | no | unknown | Brown et al. 1976 |
|  | na | **yes** | na? | testes | yes | na? | no | no | unknown | Marshall et al. 1982 |
|  | 1 | **yes** | na? | testes | right-sided abdominal | na | no | no | unknown | Schmerbach et al. 2005 |
|  | 2 | **yes** | uterine body, bilateral oviducts, cranial vagina | testes | ?? | epididymis, prostate, vas deferens, penis | no | **yes** | **exon 3 (AMHR2)** | Wu et al. 2009 |
|  | 1 | **uterus-like** | uterine horns, vagina | testes | bilateral abdominal | prostate | no | no | unknown | Matsuu et al. 2009 |
|  | 1 | ?? | uterine horns | testes | right-sided abdominal | prostate | no | no | unknown | Vegter et al. 2010 |
|  | 1 | **yes** | uterine horns, cervix | testes (atrophic) | right-sided abdominal | epididymis, vas deferens, scrotum | no | **yes** | **exon 3 (AMHR2)** | Dzimira et al. 2018 |
|  | 1 | **yes** | na | na | no | prostate, penis, scrotum with testes | no | **yes** | **exon 3 (AMHR2)** |  |
|  | 1 | **yes** | cranial vagina, bilateral oviducts | testes | no | prostate | no | **yes** | **exon 3 (AMHR2)** | Nogueira et al. 2019 |
|  | 1 | **yes** | uterine horns, oviducts, vaigna | testes | no | epididymis, prostate, scrotum, penis | no | **yes** | **exon 3 (AMHR2)** | Welsh et al 2023 |
| **Yorkshire Terrier** | 1 | **yes** | na? | testes | bilateral | na? | no | no | unknown | Hagel et al. 2010 |
|  | 1 | **yes** | vagina, vulva, cervix | gonads (resembling testes) | ?? | epididymis, rudimentary deferens vas, | no | no | unknown | Dianovský et al. 2013 |
|  | 1 | ?? | uterine horns | testes | ?? | prostatic-like structure, rudimentary penis | no | no | unknown | Silva et al. 2018 |
|  | 1 | **yes** | na | testes | no | penis | **WGS** | **WGS** | unknown | Nowacka-Woszuk et al. 2022; **this study** |
|  | 1 | **yes** | uterine horns, left-sided ovarian tissue | na | na | na | no | no | unknown | Szabo et al.. 2023 |
| **Mixed breed** | 1 | **yes** | uterine horns, cranial vagina | testes | left-sided inguinal, right-sided abdominal | malformed penis, hypospadia | no | no | unknown | Kuiper et al. 2004 |
| **Basset Hound** | 1 | **uterus-like** | uterine horns | testes? | no? | vas deferens | no | no | unknown | Pop et al. 2015 |
|  | 1 | **uterus-like?** | uterine horns? | testes? | no? | male genitalia  **(low AMH activity!)** | no | **yes** | unknown | Pop et al. 2017 |
| **German Shepherd** | 1 | **yes** | single uterine horn, cervix, blind vaigina, vulvar labia | right ovotestis | no | hypoplastic penis, one-sided epididymis, vas deferens | **yes (ex1-4)** | **yes** | unknown | De Lorenzi et al. 2018 |
| **Belgian Malinous** | 1 | **yes** | uterine horns | testes | right-sided | epididymis, penis, scrotum? | **yes (ex1-5)** | **yes** | unknown | Smit et al. 2018 |
| **Pomeranian** | 1 | **yes** | uterine horns | na | na | prostate | no | no | unknown | Vignoli et al. 2020 |
| **Cocker Spaniel** | 1 | **yes** | uterine horns | hypoplastic gonads (resembling testes) | bilateral abdominal | hypoplastic prepuce and penis, lack of scrotum | no | no | unknown | Cinti et al. 2021 |

S**upplementary Table *2*.** Summary of mapping quality for four canine genomes.

| **Genome assembly** | **Mapped reads** | **Unmapped reads** | **Duplication rate** | **GC Percentage** | **Mean coverage (standard deviation)** | **Mean mapping quality** | **General error rate** |
| --- | --- | --- | --- | --- | --- | --- | --- |
| **CanFam3.1** | 718,204,481 / 99.2% | 5,806,423 / 0.8% | 20.63% | 42.06% | 43.3619 (1,623.6498) | **19.5** | 0.6% |
| **ROS_Cfam_1.0** | 719,625,347 / 99.43% | 4,158,342 / 0.57% | 20.35% | 42.12% | 43.7601 (271.0434) | **42.68** | 0.67% |
| **CanFam4** | 719,814,029 / 99.44% | 4,073,586 / 0.56% | 20.46% | 42.12% | 42.2893 (1,244.4505) | **16.15** | 0.58% |
| **CanFam6** | 702,075,819 / 98.73% | 9,017,541 / 1.27% | 20.85% | 42.22% | 44.9575 (576.3185) | **49.53** | 0.65% |

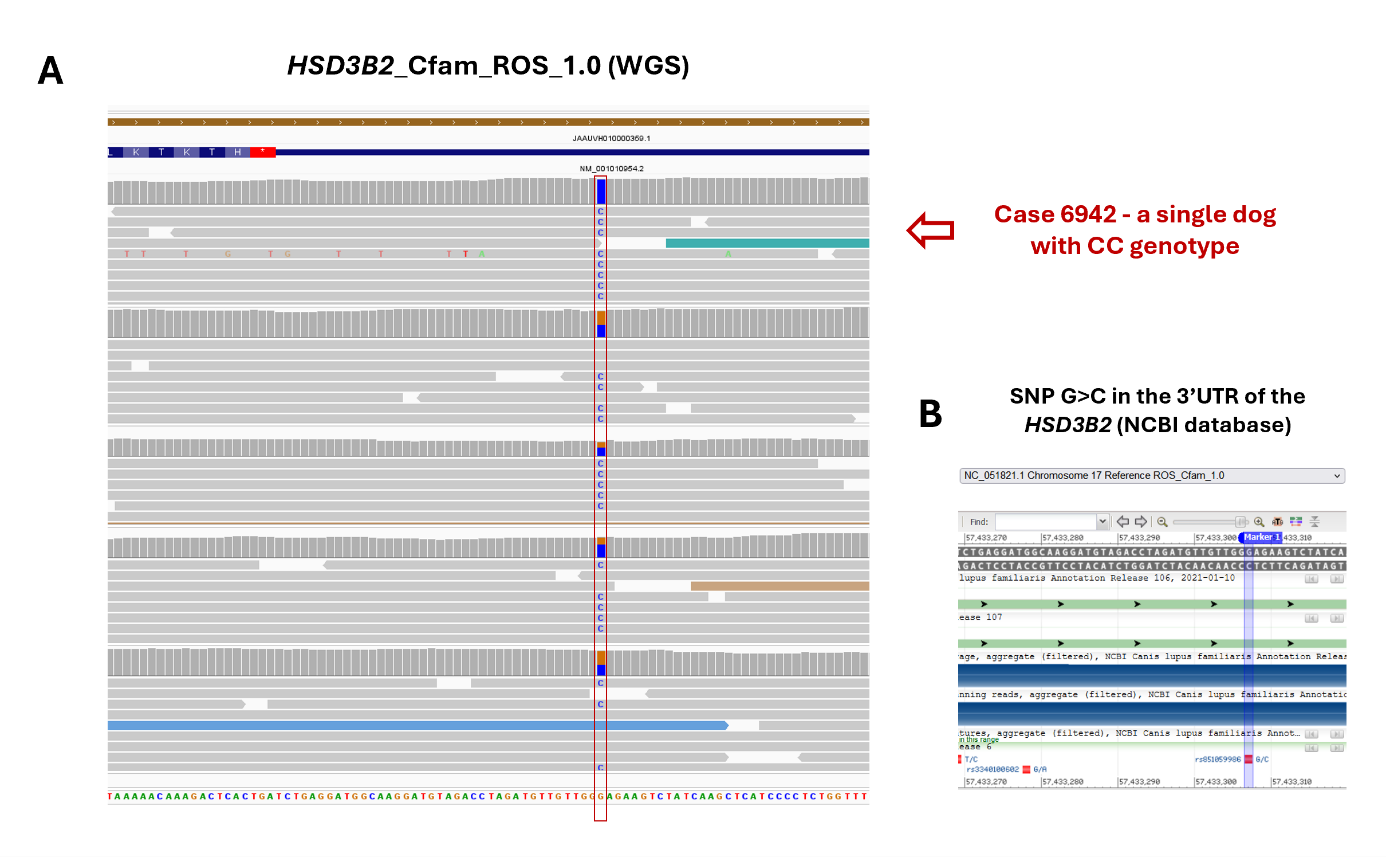

**Supplementary Figure 1.** The HSD3B2 gene was used to identify raw WGS data for a single PMDS-affected dog. **A)** WGS mapping results for five Yorkshire Terriers. Red arrow indicates PMDS-affected dog, which was the only homozygote at the indicated locus. **B)** The indicated SNP in the 3’region of the HSD3B2 gene as recorded in the NCBI database.

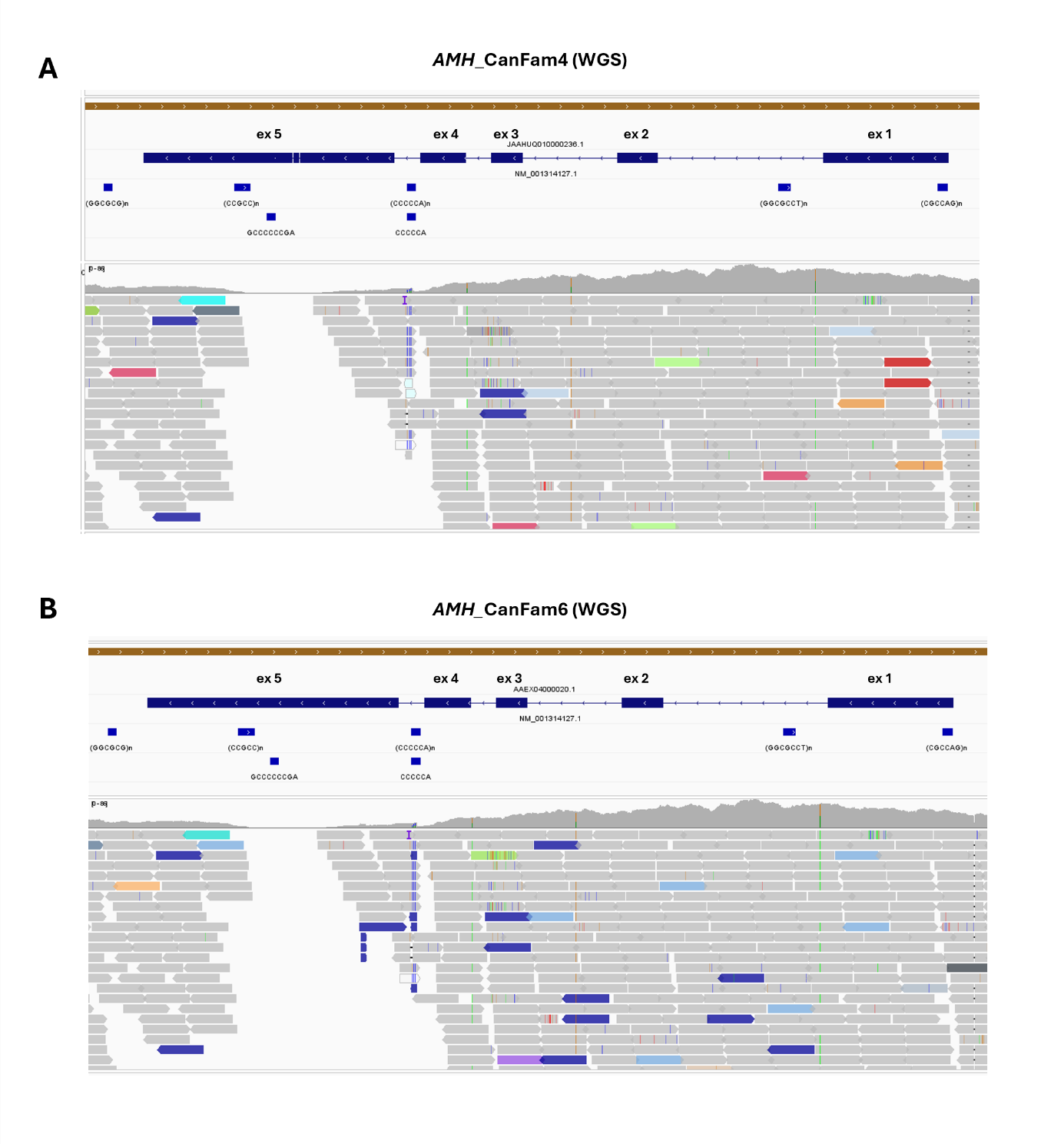

**Supplementary Figure 2.** The coverage of canine AMH gene with WGS reads mapped to **(A)** CanFam4 and **(B**) CanFam6 genome assemblies for dog #6942 diagnosed with PMDS.

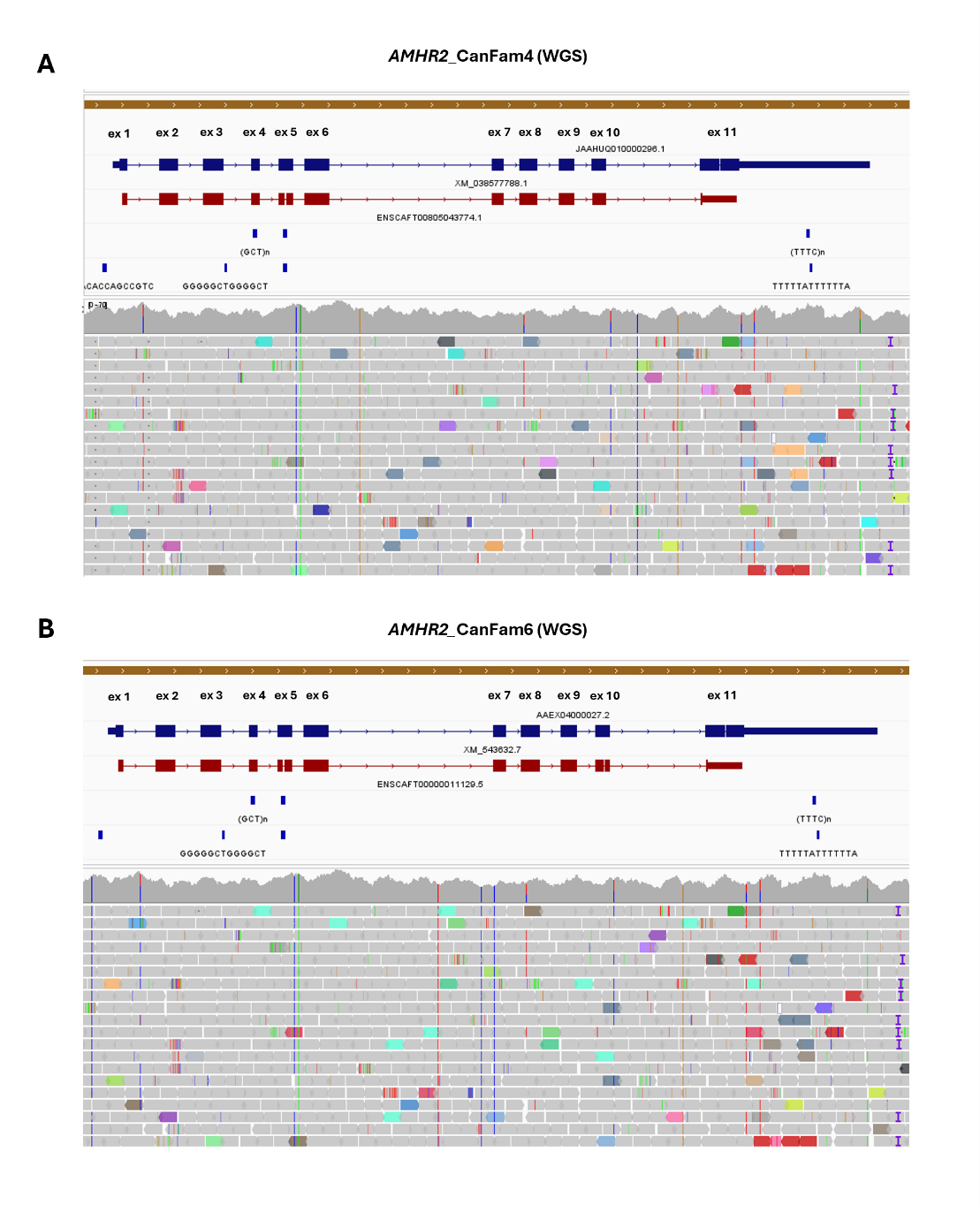

**Supplementary Figure 3.** Coverage of canine AMHR2 gene with WGS reads mapped to the CanFam4 **(A)** and CanFam6 **(B)** genome assemblies for dog #6942 diagnosed with PMDS.

**Supplementary Figure 4.** Alignment of the C-terminal part of AMHR2 protein sequences from selected species: mouse (Q8K592), dog (A0A8C0TD47), cat (M3W4X5), bovine (E1BHR7), human (Q16671) and pig (A0A4X1WCQ3), indicating that the canine and feline proteins are shorter compared to those of other species.

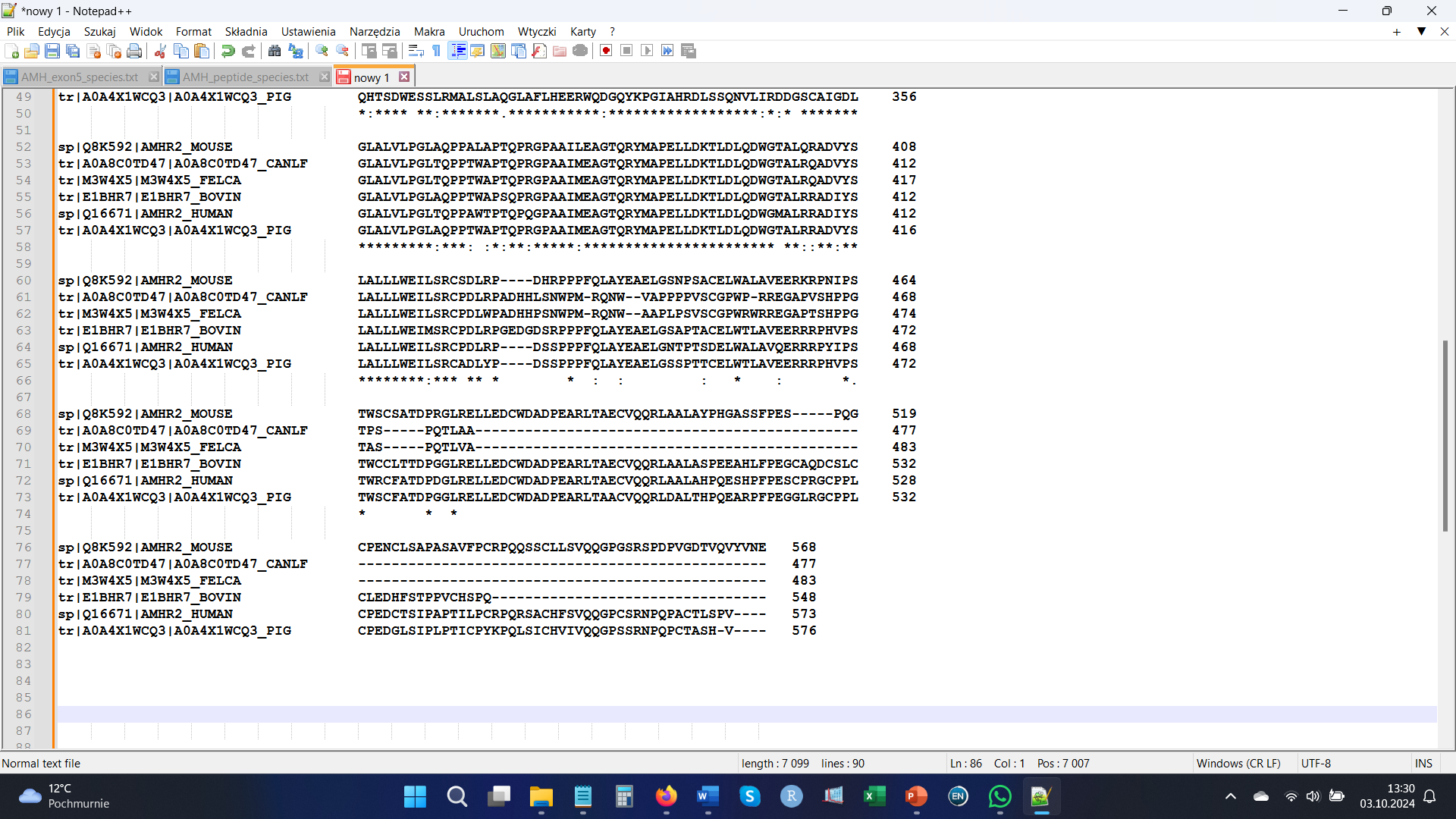

**C-terminal end of the AMHR2 proteins (based on Uniprot database)**

**Supplementary Figure 4.**

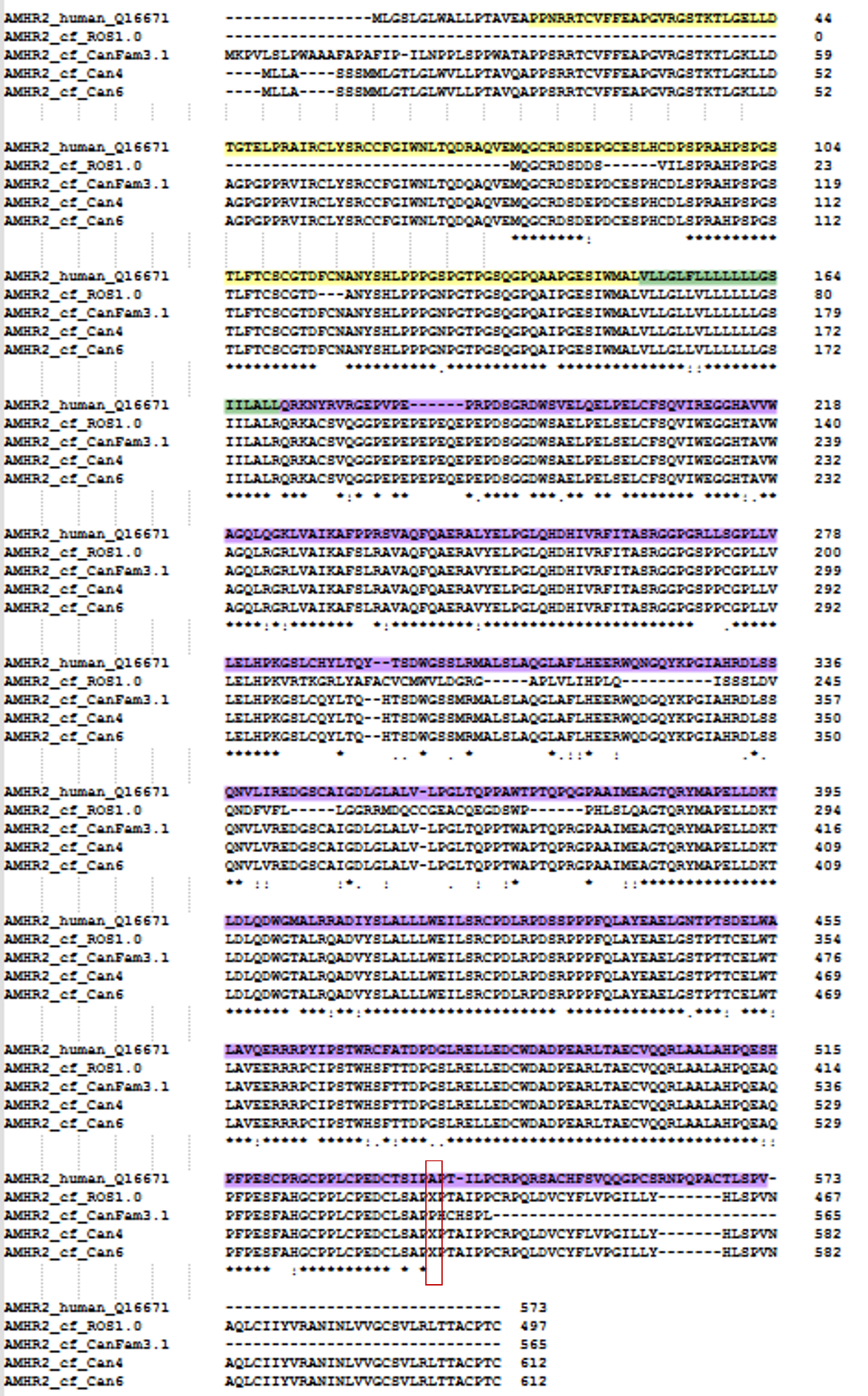

**Supplementary Figure 5.** The alignment of human and canine AMHR2 protein sequences, including four canine genome assemblies. The extracellular domain is highlighted in yellow, the transmembrane domain in green, and the cytoplasmic domain in purple. Positions with errors ('X') are marked with a red frame.

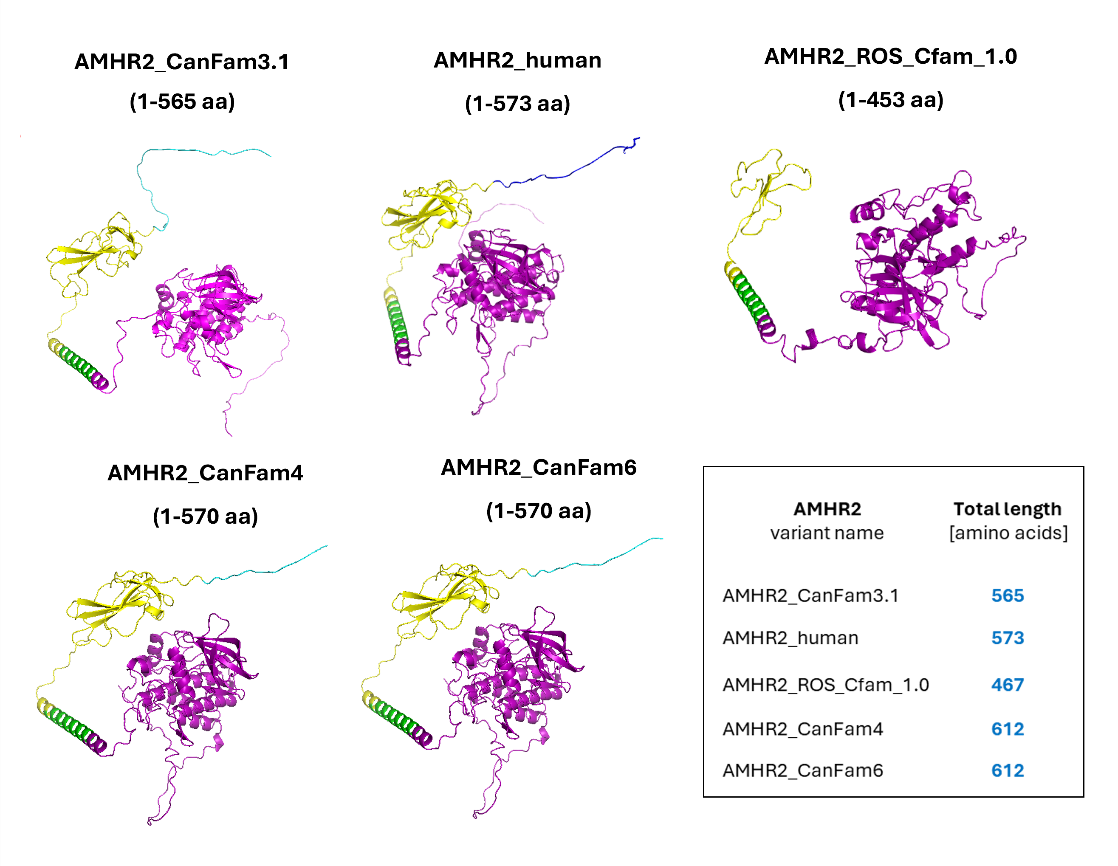

**Supplementary Figure 6.** Predicted protein structures for human and four canine AMHR2 sequences. Signal peptides are highlighted in blue, the extracellular domain in yellow, the transmembrane domain in green, and the cytoplasmic domain in purple. The lengths of the predicted structures are indicated above the figures. A frame provides information about the total protein lengths.
